# Dissociated network and cellular remodeling in the hippocampal-cortical circuit after induction of neuronal hyperexcitability

**DOI:** 10.64898/2026.11.18.767525

**Authors:** Diego Varona-de la Hera, Juan David Osejo-Ocampo, Leire Boveda-Altube, Miguel Aguilera, Juan Manuel Encinas-Perez, Nerea Llamosas

## Abstract

Neuronal hyperexcitability (NHE) can leave lasting consequences in neuronal circuits. It is often assumed that hyperexcitable neurons give rise to hyperactive networks. Here we asked how a single injection of kainic acid, which induces NHE without seizures, reshapes the hippocampal-cortical circuit at the cellular and network levels two months later. Using high-density microelectrode arrays in acute slices, we found a long-lasting reduction in spontaneous firing and bursting across the hippocampal regions, a lower proportion of active electrodes throughout the circuit, and weaker coupling from CA1 to the entorhinal cortex. In contrast, whole-cell recordings revealed region- and cell-type-specific changes in intrinsic excitability. CA1 and CA3 pyramidal neurons and CA3 interneurons exhibited increased evoked firing, whereas CA1 interneurons exhibited reduced evoked firing. Deep-layer entorhinal neurons, however, received less excitatory synaptic input and fired less. Thus, network and cellular alterations were dissociated after NHE. Finally, a pharmacological challenge with the GluK1 agonist ATPA revealed region-specific changes in network responsiveness, with a blunted response in CA3 and enhanced response in the non-entorhinal cortex, with no differences in DG, CA1 or entorhinal cortex. These findings suggest that, after the induction of NHE, region- and cell-type-specific neuronal remodeling coexists with reduced network activity ex vivo, and that the state of the circuit cannot be inferred from the properties of its individual neurons.

## Introduction

Neuronal hyperexcitability (NHE) can leave lasting consequences in neuronal circuits. It is not exclusive to epilepsy: it also occurs, often without overt seizures, in other conditions such as Alzheimer’s disease ^1–4^, Parkinson’s disease ^5^ and traumatic brain injury ^6^,where it has been proposed to contribute to circuit dysfunction ^3,7^. It is often assumed that hyperexcitable neurons give rise to hyperactive networks. Here we asked how a single injection of kainic acid, which induces NHE without seizures, reshapes both the intrinsic and synaptic properties of individual neurons and the activity of the hippocampal-cortical network two months later.

NHE triggered by an insult can outlast the insult by a long time. After experimental head trauma, for example, increased excitability of hippocampal circuits and deficits in GABAergic function persist for weeks to months^8–10^, and a single status epilepticus can produce a long-lasting change in neuronal firing mode ^11^. However, the cellular and network consequences of an insult do not necessarily go in the same direction ^12^. In organotypic hippocampal cultures, kainate has opposite regional effects, damping burst firing in CA3 while increasing excitability in CA1 ^13^, and cortical neurons can show early hypoactivity that later turns into hyperactivity ^14^. Neurons with marked synaptic alterations may even lack somatic hyperexcitability ^15^. Homeostatic mechanisms that adjust intrinsic and synaptic properties to stabilize activity ^16–19^ can explain how the properties of single cells and the activity of the network drift apart, intrinsic excitability itself is plastic^20^. Cellular and network readouts are rarely measured together in the same model, which leaves open how they relate after a hyperexcitability insult.

The hippocampal-cortical circuit is well suited to address this question. Information flows through a loop that links the entorhinal cortex (EC), the dentate gyrus (DG), CA3 and CA1, with CA1 projecting back to the deep layers of the EC ^21^. These deep layers are highly vulnerable to hyperexcitability ^22,23^, and a persistent change at any node could propagate through the loop. Assessing it therefore requires monitoring several regions simultaneously, as well as the properties of the excitatory and inhibitory neurons that compose them ^24,25^.

To do this, we used a model in which a single intrahippocampal injection of a low concentration of kainic acid induces NHE ^26,27^. In this model, hyperexcitability persists for at least 50 days, when in vivo recordings reveal epileptiform spikes ^26^, but no seizures are observed and tissue markers do not converge with those of epilepsy ^27–29^. We combined high-density microelectrode array (HD-MEA) recordings of spontaneous activity across the DG, CA3, CA1, EC and non-entorhinal cortex in acute slices ^30^ with whole-cell patch-clamp recordings from pyramidal neurons and GABAergic interneurons of CA1 and CA3 and from deep-layer EC neurons, and we pharmacologically challenged the network with the kainate-receptor agonist ATPA ^31^ to probe its responsiveness.

We found that, two months after the injection, spontaneous activity was reduced in hippocampal regions and the directed coupling from CA1 to EC was weakened, whereas single neurons showed heterogeneous changes in intrinsic and synaptic properties that did not follow the direction of the network. The response to ATPA was selectively altered in CA3 and in the non-entorhinal cortex. Together, these results show that NHE without seizures leaves a persistent, region-specific remodeling in which cellular and network properties are dissociated.

## Materials and Methods

### Animals

All experiments were performed in male and female C57BL/6 mice. Intrahippocampal kainic acid or saline injections were performed at 2 months of age, and electrophysiological recordings were conducted approximately 2 months later, at 4 months of age. Mice were maintained with ad libitum access to food and water under controlled temperature (19-22 °C) and humidity (40-50%) conditions, with a 12-h light/dark cycle. All experimental procedures were approved by the Ethics Committees of the University of the Basque Country (UPV/EHU) and the Provincial Council of Bizkaia (Bizkaiko Foru Aldundia), and were conducted in accordance with European Directive 2010/63/EU and the National Institutes of Health guidelines for the care and use of laboratory animals.

### Stereotaxic surgery and induction of neuronal hyperexcitability

#### Intrahippocampal kainic acid injections

Neuronal hyperexcitability (NHE) was induced by the administration of kainic acid (KA, Sigma, #K0250) as previously described ^26,27^. Mice were anesthetized with isoflurane (3-4 % for induction and 1-2 % for maintenance) and received a single subcutaneous dose of buprenorphine (0.1 mg/kg;) for analgesia. Animals were positioned in a stereotaxic apparatus and the right hippocampus was targeted using the following coordinates relative to bregma: anteroposterior (AP), -1.8 mm; mediolateral (ML), -1.6 mm; and dorsoventral (DV), -1.9 mm. A total volume of 50 nL of KA (0.74 mM) was injected using a glass micropipette coupled to a nanoinjector (Nanoject II, Drummond Scientific, USA). After injection, the micropipette was left in place for 2 minutes to prevent reflux and then slowly withdrawn. Control animals underwent the same surgical procedure and received an equivalent volume of Sterile 0.9% NaCl solution (physiological saline). Following surgery, the incision was sutured and animals were allowed to recover on a heating pad.

#### Intrahippocampal viral-vector injection

To enable targeted whole-cell patch-clamp recordings from GABAergic interneurons in the CA1 and CA3 hippocampal regions, mice received a single stereotaxic injection of an adeno-associated viral vector expressing GFP under the control of the mDLX regulatory element (AAV9-mDLX-GFP). The virus was injected 10 to 15 days before electrophysiological recordings. Using the stereotaxic procedure described above, a glass micropipette was positioned at the following coordinates relative to bregma: anteroposterior (AP), −1.8 mm; mediolateral (ML), −1.6 mm; and dorsoventral (DV), −1.9 mm. A total volume of 500 nL was injected over 5 minutes using a Nanoject II nanoinjector. The micropipette was left in place for an additional 5 minutes before being slowly withdrawn. GFP-positive interneurons were subsequently identified in acute hippocampal slices for whole-cell patch-clamp recordings.

### Acute brain slice preparation

Acute brain slices from the ipsilateral hemisphere were prepared as previously described ^32–35^ for whole-cell patch-clamp and high-density microelectrode array (HD-MEA) recordings using a common procedure, with differences in slice orientation and artificial cerebrospinal fluid (aCSF) composition depending on the experimental approach.

Four-month-old mice of either sex were anesthetized with isoflurane and decapitated in accordance with approved animal experimental procedures. Brains were rapidly removed and immersed in ice-cold cutting solution containing (in mM): 92 NMDG, 2.5 KCl, 1.25 NaH₂PO₄, 30 NaHCO₃, 20 HEPES, 25 glucose, 2 thiourea, 5 sodium ascorbate, 3 sodium pyruvate, 0.5 CaCl₂·4H₂O, and 10 MgSO₄ (pH adjusted to 7.3-7.4 with 4 N HCl).

Sagittal slices (300 μm) were prepared for whole-cell patch-clamp recordings, whereas horizontal slices (300 μm) were obtained for HD-MEA recordings using a vibrating microtome (Leica Biosystems). Following sectioning, slices were transferred to a submersion chamber containing the corresponding aCSF, maintained at 32 °C for 30 min, and subsequently kept at room temperature until recording (1-6 h). 3-5 slices were obtained per animal.

For patch-clamp experiments, the aCSF contained (in mM): 124 NaCl, 2.5 KCl, 1.3 MgSO₄, 2.5 CaCl₂, 1 NaH₂PO₄, 10 D-glucose, and 26 NaHCO₃. For HD-MEA experiments, the aCSF contained (in mM): 126 NaCl, 3.5 KCl, 1.25 MgSO₄, 1.6 CaCl₂, 1.25 NaH₂PO₄, 10 D-glucose, and 26 NaHCO₃. Both solutions were continuously equilibrated with carbogen (95% O₂/5% CO₂) and maintained at pH 7.4 and 300-310 mOsm.

### Whole-cell patch-clamp recordings

After 1-1.5 h individual slices, containing the dorso-medial hippocampus and the medial entorhinal cortex were placed in the recording chamber mounted on the stage of Scientifica microscope and superfused at 2 ml/min with warm (32 ± 0.5°C). Hippocampal pyramidal neurons were identified based on their anatomical location and somatic morphology, whereas GABAergic interneurons were identified by GFP fluorescence following AAV9-mDLX-GFP injection ^36^. In the EC, putative pyramidal neurons were selected based on their location within layers V-VI, somatic morphology and electrophysiological characteristics consistent with pyramidal neurons, including relatively low firing frequencies in response to depolarizing current injections. Experiments were made with borosilicate patch pipettes (5 to 7 mΩ) filled with a solution containing the following (in mM): 130 K-gluconate, 20 KCl, 0.2 EGTA, 10 Hepes, 4 Mg-ATP, 0.3 Na-GTP (pH: 7.3, 280 mOsm). Series resistance was monitored without compensation with 5mV depolarizing steps (200ms) applied every 60 s to ensure sufficient and stable electrical access to the cell. Series resistance was not compensated. Electrophysiological signals were amplified using a MultiClamp 700B amplifier (Molecular Devices), low-pass filtered at 4 kHz in voltage-clamp mode and 2 kHz in current-clamp mode, and digitized at a sampling rate of 10 kHz. Data were acquired using Clampex 10.5 and analyzed offline using Clampfit 11.3 (Molecular Devices).

Cells with an access resistance >25 MΩ or exhibiting unstable access resistance (>20% change during recording) were excluded from further analysis. Resting membrane potential was measured immediately after establishing the whole-cell configuration in current-clamp mode at zero holding current (I = 0). Membrane capacitance was calculated from the integral of the capacitive transient evoked by a 5-mV voltage step, and input resistance was determined from the corresponding steady-state current response. For voltage-clamp recordings, CA1 and CA3 pyramidal neurons were held at −70 mV, whereas inhibitory interneurons were held at −60 mV. To assess Kir- and HCN-associated currents, hyperpolarizing voltage steps from −60 to −130 mV were applied in 10-mV decrements. Rheobase was defined as the minimum depolarizing current required to evoke an action potential. The properties of the first evoked action potential were then characterized, including threshold, amplitude, half-width, rise time, decay time, and fast afterhyperpolarization (fAHP) amplitude. AP onset was defined as the point at which the first derivative of the membrane potential (dV/dt) exceeded 5 mV/ms. Neuronal firing was assessed by counting the number of APs elicited during depolarizing current steps applied in 20 pA increments. Kir currents were quantified as the difference between the current measured at the end of each voltage step and the baseline holding current at −70 mV. HCN-associated currents were quantified as the difference between the current measured at the end of each hyperpolarizing step and the initial peak current following the capacitive transient. Spontaneous postsynaptic currents were recorded in voltage-clamp mode at a holding potential of −70 mV in CA1 and CA3 pyramidal neurons and entorhinal cortex neurons, and at −60 mV in hippocampal GABAergic interneurons. Spontaneous excitatory postsynaptic currents of entorhinal cortex neurons (sEPSCs) were analyzed for event frequency, amplitude, rise time, decay time, and half-width. Individual events were identified using the semi-automated sliding-template detection algorithm implemented in Clampfit (pCLAMP 11.3; Molecular Devices). Detection templates were generated by averaging representative spontaneous events, and the detection threshold was set to 4. Only single-peak events were included in the analysis.

### High-density microelectrode array recordings

Extracellular neuronal activity was recorded from acute horizontal brain slices (300 μm) using a BioCAM DupleX high-density microelectrode array (HD-MEA) system equipped with 4096 recording electrodes (3Brain, Switzerland). Slices were continuously superfused with oxygenated artificial cerebrospinal fluid (aCSF) at a flow rate of 4 mL/min and maintained at 35-38 °C. Spontaneous extracellular activity was acquired at a sampling rate of 19.75 KHz in 3-min recording blocks across the hippocampal-cortical circuit, including the dentate gyrus (DG), CA3, CA1, entorhinal cortex (EC), and non-entorhinal cortical regions. Anatomical segmentation was performed based on microscopic images of the brain slices, allowing the identification and delineation of individual brain regions according to their anatomical landmarks.

Slices were allowed to stabilize on the chip for 10 min, after which spontaneous extracellular activity was recorded for 3 min (baseline). Slices were then superfused with aCSF containing 10 µM ATPA, a GluK1-preferring kainate receptor agonist. The effect of ATPA was assessed in a 3-min recording obtained 15 min after the onset of perfusion.

### HD-MEA data processing and network analysis

Raw extracellular signals were recorded with BrainWave 6 (3Brain) at 19.75 kHz and stored in .brw files read directly with custom Python scripts. Spikes were detected with the Precise Time Spike Detection (PTSD) algorithm ^37^. Signals were band-pass filtered between 300 Hz and 3kHz (4th-order Butterworth filter, zero-phase). A spike was detected when the peak-to-peak amplitude of the filtered signal within a peak lifetime period of 2 ms exceeded a differential threshold equal to seven times the noise standard deviation (sigma) of each electrode, and a refractory period of 1 ms was applied. The noise standard deviation is electrode-specific and was taken from the last noise measurement stored in each file; only the multiplication factor (7) was set. For each spike, the waveform was extracted from 0.5 ms before to 1.5 ms after the peak.

To asses detection threshold, we compared the noise standard deviation used for spike detection, the last noise standard deviation measurement stored in each file. For each recording, this value was averaged across all electrodes within the analyzed regions of interest.

Only recording channels with a mean firing rate between 0.02-15 spikes/s were included for the analysis, to exclude inactive channels and noise artefacts.

Bursts were defined as sequences of at least two spikes with a maximum interspike interval of 30 ms. Spontaneous neuronal activity was characterized by mean firing frequency and burst frequency, and interburst intervals (IBIs).

Active electrodes were defined as those exhibiting a spike frequency between 0.02-15 spikes/s. The percentage of active electrodes was calculated as the number of active electrodes divided by the total number of recording electrodes within each region, multiplied by 100.

To characterize differences in interregional functional organization across anatomical regions, we performed the following analysis: Spikes were detected from raw .bxr recordings using the PTSD algorithm (3Brain BrainWave), with a threshold of 7 standard deviations and a 300-3000 Hz bandpass filter. Channels with a mean firing rate below 0.05 spikes/second were discarded, and recordings retaining fewer than 50 active channels after this step were excluded from the analysis. To extract functional connectivity, spike trains were binned at 10 ms resolution and z-scored per channel. Directed functional connectivity between channel pairs was computed as a one-bin-lagged cross-correlation,

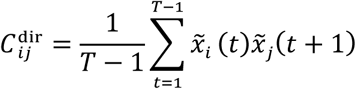

Channels were assigned to one of five hippocampal regions of interest (DG, CA3, CA1, Cortex, EC) based on their electrode position. For each ordered region pair (A, B), inter-regional connectivity (mean connectivity from region A to region B) and intra-regional connectivity (mean connectivity within region A) were computed from the directed connectivity matrix, and the relative directed connectivity strength was defined as

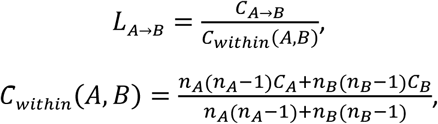

with *n_A_*, *n_B_* the channel counts per region, where *L_A_*_→*B*_ was computed only when the pooled within-region baseline was positive. Recordings with negligible net directionality across the full connectivity matrix 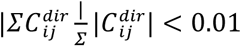 were excluded from directed-metric analysis, since in these cases the directed signal is not distinguishable from noise.

### Statistical analysis

Statistical analyses were performed using GraphPad Prism (version 9). Data normality was assessed using the D’Agostino-Pearson omnibus normality test. Comparisons between two groups were performed using Student’s t-test for normally distributed data or the Mann-Whitney U test for non-normally distributed data. Analyses involving two factors were performed using two-way ANOVA followed by appropriate post hoc multiple-comparison tests. Mixed-effects models were used for repeated-measures analyses when missing values were present. To determine whether NHE differentially affected the firing responses of pyramidal neurons and interneurons, a factorial analysis including cell type, treatment group and injected current was performed on the original firing data. Differences between NHE and SAL (Δ spikes) were calculated for graphical representation. Cumulative distributions were compared using the Kolmogorov-Smirnov test. Functional connectivity analysis was restricted to the four core hippocampal-cortical segments of the circuit (EC→DG, DG→CA3, CA3→CA1, CA1→EC), comparing SAL and NHE slices. Within each segment and group, values outside 1.5 times the interquartile range were removed before plotting and statistical analyses, as near-zero connectivity values can produce artificially large *L_A_*_→*B*_ ratios. The statistical tests applied to individual datasets are specified in the corresponding figure legends. Statistical significance was set at p < 0.05. Data are presented as mean ± SEM or median with interquartile range (IQR), as appropriate. Sample sizes are indicated in the figure legends.

## Results

### Region-dependent hypoactivity in the hippocampal-cortical circuit

We first sought to determine whether neuronal hyperexcitability (NHE), induced by a single intrahippocampal injection of a low concentration of kainic acid, leads to lasting alterations in hippocampocortical network activity. We recorded spontaneous network activity ex vivo with a high-density microelectrode array (HD-MEA, 4096 electrodes) in acute horizontal slices that preserve the hippocampal-parahippocampal circuit, 50-70 days after the injection. Electrodes were assigned to DG, CA3, CA1, medial entorhinal cortex (EC) and non-entorhinal cortex (NEC) using anatomical landmarks (Fig. 1A, B).

**Figure 1.**
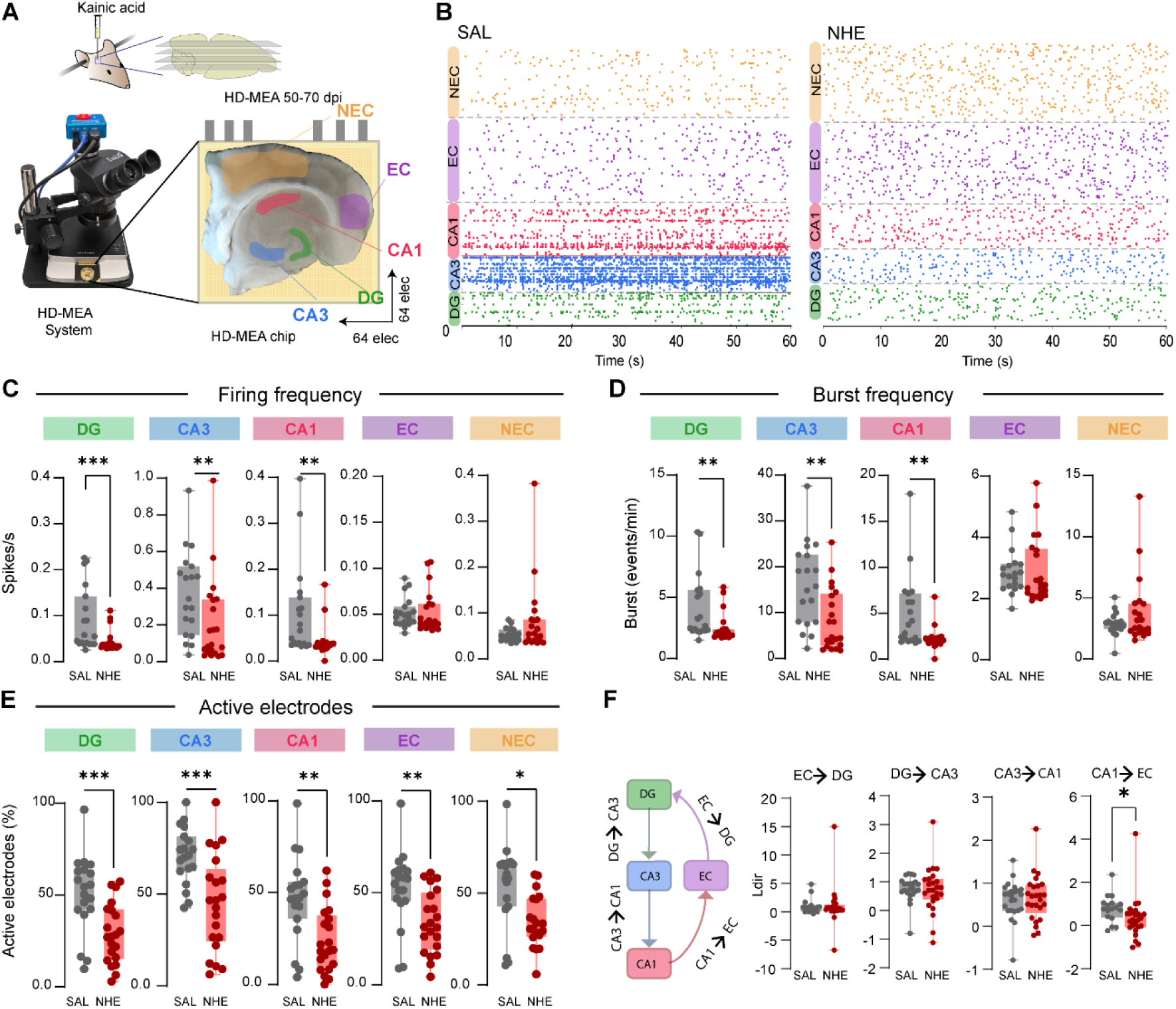
Region-dependent hypoactivity in the hippocampal-cortical circuit. (A) Experimental design. Mice received a single intrahippocampal injection of kainic acid (NHE) or saline (SAL), and acute horizontal slices were recorded on a high-density microelectrode array (HD-MEA) 50-70 days post-injection (dpi). Recording areas were delineated into the dentate gyrus (DG), CA3, CA1, entorhinal cortex (EC) and non-entorhinal cortex (NEC). (B) Raster plots of 60 s of spontaneous activity from representative SAL (left) and NHE (right) slices, color-coded by region. (C-E) Spontaneous firing frequency (C), burst frequency (D) and percentage of active electrodes (E) in each region in SAL and NHE slices. (F) Left, schematic of the hippocampal-entorhinal loop with the four directed interactions analyzed (EC → DG, DG → CA3, CA3 → CA1, CA1 → EC). Right, directional coupling index (Ldir) for each interaction. For C-E, DG, CA3, CA1 and EC: SAL n = 19 slices, NHE n = 21 slices; NEC: SAL n = 19 slices, NHE n = 20 slices, from 5 mice per group. For F, SAL n = 18 slices from 8 mice, NHE n = 21 slices from 8 mice. Mann-Whitney test for C-F. Data are shown as box-and-whisker plots (median, interquartile range and minimum-maximum), with individual slices overlaid. *p < 0.05, **p < 0.01, ***p < 0.001.

Analysis of spontaneous activity revealed region-dependent alterations in network function at 50-70 dpi. Spontaneous firing rates were significantly reduced in DG, CA3, and CA1 in the NHE group compared with saline controls, whereas no significant differences were detected in EC or NEC (Fig. 1C). Similarly, spontaneous burst frequency was significantly decreased across all three hippocampal regions, with no significant differences observed in either cortical region (Fig. 1D).

Analysis of the proportion of active electrodes revealed a more widespread alteration. This parameter was significantly reduced in the NHE group across all five recording regions, including DG, CA3, CA1, EC, and NEC (Fig. 1E). To assess whether these differences could reflect technical or signal-related factors, we compared baseline noise and spike amplitude between groups. Baseline noise (Fig. S1A, C) and the peak-to-peak amplitude of detected spikes in each region (Fig. S1B) were comparable between NHE and saline slices, indicating similar recording quality across groups and that NHE did not alter background noise or the extracellular spike signal itself. Thus, the observed differences reflect changes in the level of spontaneous activity rather than in signal properties or detectability. Together, these findings indicate that NHE is associated with a reduction in spontaneous hippocampal firing and bursting activity, accompanied by a broader decrease in the proportion of active recording electrodes throughout the hippocampal-cortical circuit.

To investigate whether these alterations in spontaneous network activity were accompanied by changes in interregional functional organization, we examined directed interactions between hippocampal and entorhinal regions using a directional coupling index (Fig. 1F). We focused on the EC → DG, DG → CA3, CA3 → CA1, and CA1 → EC interactions, representing a sequence of directed functional relationships within the hippocampal-cortical circuit. While directional coupling remained comparable between NHE and saline groups for EC → DG, DG → CA3, and CA3 → CA1, a significant reduction was observed in the CA1 → EC interaction in NHE mice (Fig. 1F).

Collectively, these findings reveal alterations in hippocampal-cortical network organization two months after NHE, characterized by reduced spontaneous activity in hippocampal regions, a widespread decrease in the proportion of active electrodes in hippocampus and cortex, and a selective reduction in directed functional coupling from CA1 to EC.

### Divergent remodeling of intrinsic excitability in CA1 pyramidal neurons and interneurons

Given the reduced spontaneous firing and bursting activity, together with the lower proportion of active electrodes observed in CA1 using HD-MEA recordings, we next investigated whether these network-level alterations were accompanied by changes in the intrinsic excitability of CA1 excitatory and inhibitory neurons. To address this question, we performed whole-cell patch-clamp recordings from CA1 pyramidal neurons and inhibitory interneurons in acute hippocampal slices obtained from control and NHE mice (Fig. 2A, J).

**Figure 2.**
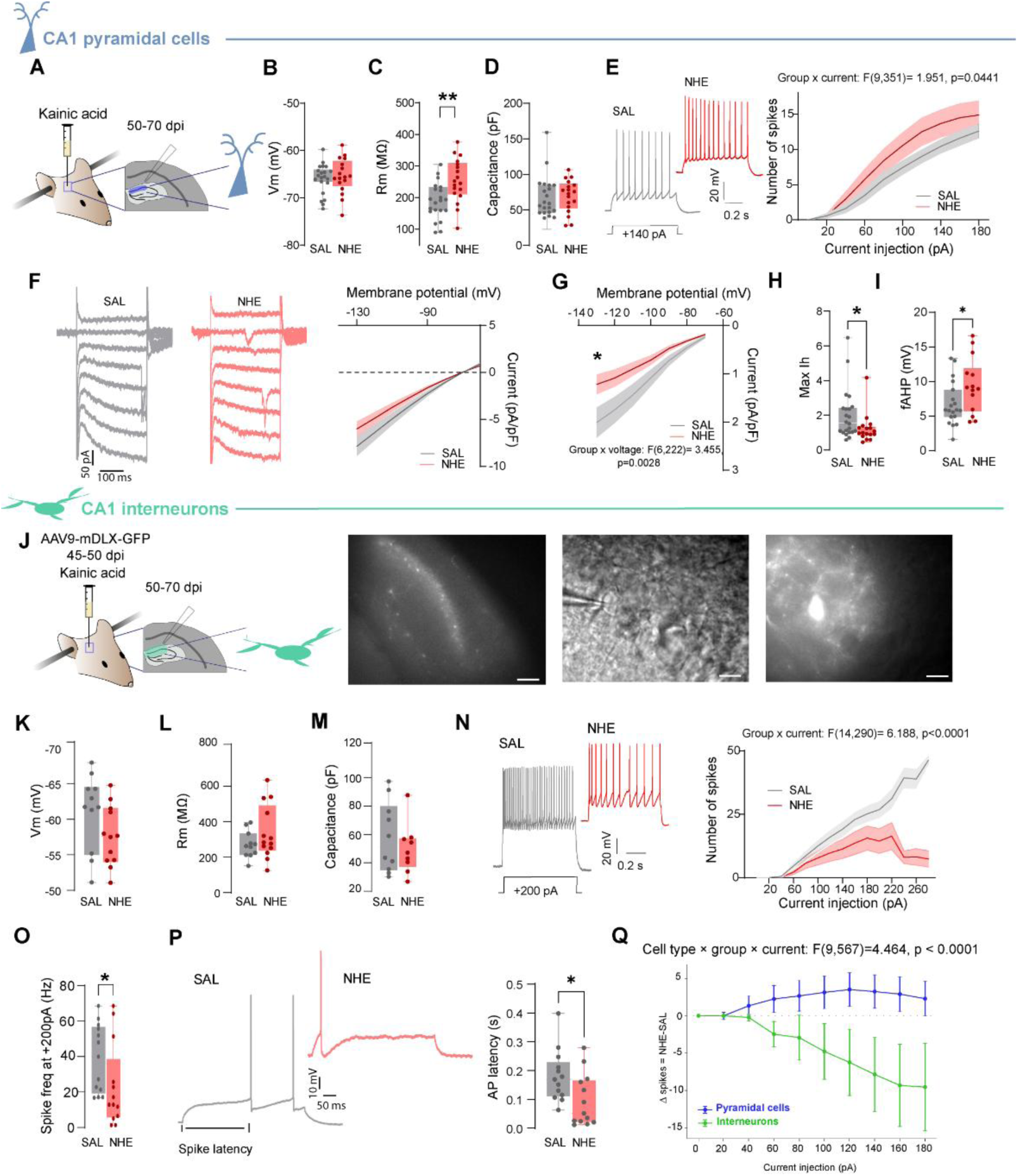
Divergent remodeling of intrinsic excitability in CA1 pyramidal neurons and interneurons. (A) Experimental design for whole-cell patch-clamp recordings of CA1 pyramidal neurons in acute sagittal slices from SAL and NHE mice at 50-70 days post-injection (dpi). (B-D) Resting membrane potential (Vm; B), input resistance (Rm; C) and membrane capacitance (D) of CA1 pyramidal neurons. (E) Left, representative voltage responses to a depolarizing current step (+140 pA). Right, input-output curves showing the number of action potentials evoked by increasing current injections (group × current interaction: F(9,351) = 1.951, p = 0.0441). (F) Representative voltage-clamp currents evoked by hyperpolarizing voltage steps (left) and current-voltage relationship of the steady-state inward current (right). (G) Current-voltage relationship of the hyperpolarization-activated inward current (group × voltage interaction: F(6,222) = 3.455, p = 0.0028). (H) Maximal hyperpolarization-activated inward current. (I) Fast afterhyperpolarization (fAHP) amplitude. (J) Experimental design for recordings from CA1 interneurons, identified by AAV9-mDLX-GFP expression (virus injected at 45-50 dpi), and representative fluorescence image of CA1 (left), DIC image with the recording pipette (middle) and fluorescence image of a GFP-positive interneuron (right). (K-M) Vm (K), Rm (L) and capacitance (M) of CA1 interneurons. (N) Left, representative voltage responses to a depolarizing current step (+200 pA). Right, input-output curves of CA1 interneurons (group × current interaction: F(14,290) = 6.188, p < 0.0001). (O) Action potential firing frequency at +200 pA. (P) Representative traces illustrating the latency to the first action potential and its quantification. (Q) Difference in the number of action potentials between NHE and SAL neurons (Δ spikes = NHE − SAL) across current steps for pyramidal neurons (blue) and interneurons (green) (cell type × group × current interaction: F(9,567) = 4.464, p < 0.0001). Pyramidal neurons: SAL n = 23 cells from 10 mice, NHE n = 18 cells from 6 mice; interneurons: SAL n = 14 cells from 6 mice, NHE n = 13 cells from 7 mice. Unpaired t-test or Mann-Whitney test for B-D, H, I, K-M, O and P. Two-way repeated-measures ANOVA or mixed-effects model for E, G and N. For Q, a three-factor analysis was performed on the original firing data, with cell type, treatment group and injected current as factors. Data are shown as box-and-whisker plots (median, interquartile range and minimum-maximum) with individual cells overlaid, and as mean ± SEM (shaded) for current-response and current-voltage curves. *p < 0.05, **p < 0.01. Scale bars: 100 µm (J, left), 20 µm (J, middle), 20 µm (J, right).

We first examined the intrinsic membrane properties and firing responses of CA1 pyramidal neurons. While resting membrane potential and membrane capacitance remained comparable between groups (Fig. 2B,D), input resistance was significantly increased in NHE neurons (Fig. 2C). Consistent with an altered intrinsic input-output relationship, NHE pyramidal neurons generated more action potentials in response to depolarizing current injections, as revealed by a significant group × current interaction (Fig. 2E). Analysis of hyperpolarization-evoked currents revealed no alterations in steady-state inward currents (Fig. 2F). However, Ih-like currents, quantified as the difference between the initial and steady-state currents, were reduced in NHE pyramidal neurons (Fig. 2G, H). In addition, NHE pyramidal neurons exhibited an increased fast afterhyperpolarization (fAHP) amplitude of the action potential (Fig. 2I). Together, these findings indicate that NHE induces a long-term increase in the intrinsic excitability of CA1 pyramidal neurons, characterized by enhanced action potential firing, increased input resistance, reduced Ih-like currents, and altered action potential afterhyperpolarization.

We next investigated whether CA1 interneurons underwent similar functional changes. To identify inhibitory neurons, we used AAV9-mDLX-GFP labeling and performed whole-cell recordings from GFP-positive cells in the CA1 stratum pyramidale (Fig. 2J). Resting membrane potential, input resistance, and membrane capacitance did not show significant differences between control and NHE interneurons (Fig. 2K-M). Contrary to pyramidal neurons, interneurons from NHE mice exhibited a marked reduction in action potential firing in response to depolarizing current injections, with a significant group × current interaction (Fig. 2N). This reduction was also evident in the firing frequency measured at +200 pA (Fig. 2O). Moreover, NHE interneurons displayed a shorter latency to the first action potential (Fig. 2P), indicating that the reduction in repetitive firing was accompanied by a distinct alteration in spike initiation dynamics. Together, these findings indicate that pathological hyperexcitability induces a long-term reduction in the repetitive firing capacity of CA1 inhibitory interneurons, despite a shorter latency to action potential initiation.

Given the complementary roles of pyramidal neurons and inhibitory interneurons in regulating hippocampal circuit activity, we next sought to determine whether NHE differentially affects their evoked firing within CA1. To this end, we calculated the difference in the number of action potentials between NHE and control neurons at each injected current step (Fig. 2Q). Positive values indicate increased firing in NHE neurons, whereas negative values indicate reduced firing. This comparison revealed a significant cell type × group × current interaction, reflecting opposite responses in the two populations: pyramidal neurons exhibited a modest increase in evoked firing, whereas interneurons showed a progressive reduction from ∼60 pA onward, resulting in a larger magnitude of change in interneurons (Fig. 2Q).

Collectively, these findings demonstrate that NHE induces opposing long-term changes in the firing capacity of excitatory and inhibitory neurons within CA1. Importantly, these alterations coexist with the reduced spontaneous firing and bursting activity observed at the network level, highlighting a divergence between intrinsic neuronal excitability and spontaneous circuit activity. Having identified opposing changes in the firing responses of excitatory and inhibitory neurons in CA1, we next investigated whether similar alterations in intrinsic excitability occurred in CA3, another key region of the hippocampal circuit that also exhibited significantly reduced spontaneous firing and bursting activity in HD-MEA recordings (Fig. 1C, D).

### Increased evoked firing of CA3 pyramidal neurons and interneurons

Having identified opposing changes in the firing responses of excitatory and inhibitory neurons in CA1, we next investigated whether long-term alterations in intrinsic excitability also occurred in CA3, another key region of the hippocampal circuit that exhibited reduced spontaneous firing and bursting activity in HD-MEA recordings (Fig. 1C, D).

We first explored the intrinsic properties of CA3 pyramidal neurons. Resting membrane potential, input resistance, and membrane capacitance were comparable between groups (Fig. 3A-C). However, NHE neurons exhibited increased action potential firing at higher depolarizing current intensities, supported by a significant group × current interaction (Fig. 3D). Interestingly, this enhanced firing response was accompanied by an increased rheobase and a longer latency to action potential initiation (Fig. 3E-G), indicating that NHE neurons required stronger depolarizing currents to initiate firing but exhibited enhanced repetitive firing at higher current intensities. In addition, NHE neurons exhibited significant changes in action potential waveform properties, including increased amplitude, reduced half-width, shorter rise and decay times, and enhanced fast afterhyperpolarization (fAHP; Fig. 3H-J and Fig. S2A-C). In contrast, the hyperpolarization-evoked currents examined did not show significant overall differences between groups (Fig. S2D-E). Together, these findings reveal a complex pattern of long-term intrinsic electrophysiological remodeling in CA3 pyramidal neurons, characterized by reduced responsiveness at action potential initiation, enhanced repetitive firing during stronger depolarization, and action potential waveform remodeling.

**Figure 3.**
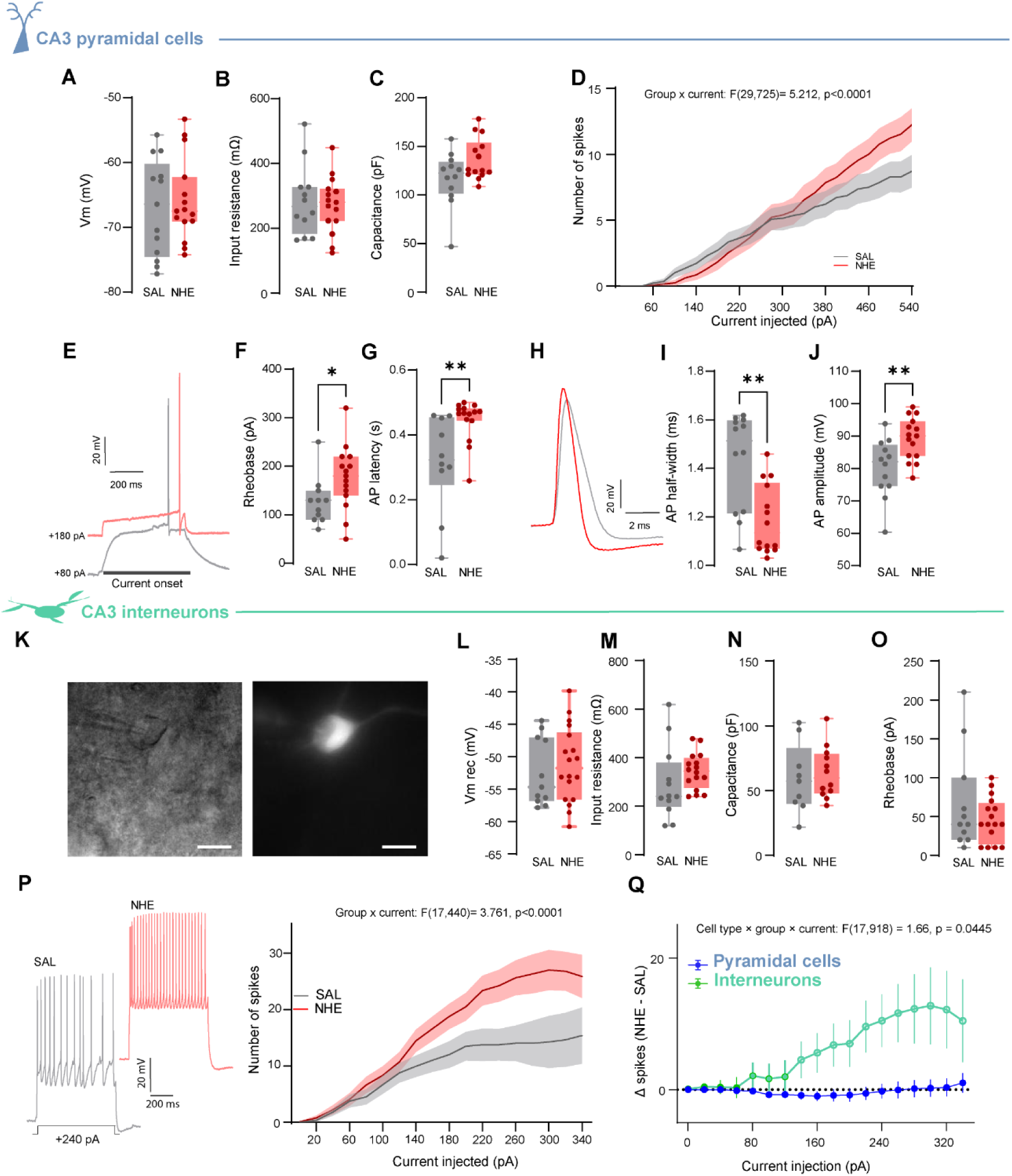
Increased evoked firing of CA3 pyramidal neurons and interneurons. (A-C) Resting membrane potential (Vm; A), input resistance (Rm; B) and membrane capacitance (C) of CA3 pyramidal neurons from SAL and NHE mice. (D) Input-output curves showing the number of action potentials evoked by increasing current injections in CA3 pyramidal neurons (group × current interaction: F(29,725) = 5.212, p < 0.0001). (E) Representative voltage responses of CA3 pyramidal neurons at near-rheobase current injection (SAL, +80 pA; NHE, +180 pA). (F, G) Rheobase (F) and latency to the first action potential (G). (H) Representative action potential waveforms. (I, J) Action potential half-width (I) and amplitude (J). (K) Representative DIC (left) and fluorescence (right) images of a CA3 interneuron identified by AAV9-mDLX-GFP expression. (L-O) Vm (L), Rm (M), capacitance (N) and rheobase (O) of CA3 interneurons. (P) Left, representative voltage responses of CA3 interneurons to a depolarizing current step (+240 pA). Right, input-output curves of CA3 interneurons (group × current interaction: F(17,440) = 3.761, p < 0.0001). (Q) Difference in the number of action potentials between NHE and SAL neurons (Δ spikes = NHE − SAL) across current steps for pyramidal neurons (blue) and interneurons (green) (cell type × group × current interaction: F(17,918) = 1.66, p = 0.0445). Pyramidal neurons: SAL n = 13 cells from 5 mice, NHE n = 13 cells from 5 mice; interneurons: SAL n = 12 cells from 6 mice, NHE n = 18 cells from 9 mice. Unpaired t-test or Mann-Whitney test for A-C, F, G, I, J and L-O; two-way repeated-measures ANOVA or mixed-effects model for D, P and Q. Data are shown as box-and-whisker plots (median, interquartile range and minimum-maximum) with individual cells overlaid, and as mean ± SEM (shaded) for input-output curves. *p < 0.05, **p < 0.01. Scale bars: 20 µm (K).

We next examined CA3 inhibitory interneurons identified by AAV9-mDLX-GFP expression (Fig. 3K). Resting membrane potential, input resistance, membrane capacitance, and rheobase did not differ significantly between control and NHE mice (Fig. 3L-O). In contrast, CA3 interneurons from NHE mice generated more action potentials in response to depolarizing current injections, supported by a significant group × current interaction (Fig. 3P). This enhanced firing response occurred without significant changes in action potential waveform properties, latency, or fAHP (Fig. S2F-K). However, analysis of hyperpolarization-evoked currents revealed significant group × voltage interactions in Kir- and HCN-associated current responses (Fig. S2L-M). Together, these findings indicate that NHE enhances the repetitive firing capacity of CA3 interneurons without detectable alterations in passive membrane properties or individual action potential waveform parameters, while also modifying their voltage-dependent membrane current responses.

As in CA1, we next determined whether NHE differentially affects the evoked firing of pyramidal neurons and interneurons within CA3, by calculating the difference in the number of action potentials between NHE and control neurons at each injected current step (Fig. 3Q). This comparison revealed a significant cell type × group × current interaction, reflecting divergent responses in the two populations: interneurons showed a marked increase in evoked firing from ∼140 pA onward, whereas pyramidal neurons showed minimal changes within the same current range, resulting in a larger magnitude of change in interneurons (Fig. 3Q). In pyramidal neurons, increased evoked firing became apparent only at higher current intensities (Fig. 3D).

Collectively, these findings reveal a distinct pattern of long-term electrophysiological remodeling in CA3 compared with CA1, with enhanced repetitive firing responses in both pyramidal neurons and inhibitory interneurons rather than the opposing changes observed in CA1. Notably, these increases in evoked firing capacity coexist with reduced spontaneous firing and bursting activity at the network level, further highlighting the divergence between intrinsic neuronal excitability and spontaneous hippocampal network activity following NHE.

### Deep-layer entorhinal neurons show reduced evoked firing and excitatory synaptic input

Given the reduced proportion of active electrodes in the entorhinal cortex (EC), despite unchanged spontaneous firing and bursting frequency, together with the selective reduction in directed coupling from CA1 to EC, we next investigated whether these network-level findings were accompanied by alterations in the intrinsic and synaptic properties of EC neurons. To this end, we performed whole-cell patch-clamp recordings from putative pyramidal neurons located in layers V-VI of the EC (Fig. 4A).

**Figure 4.**
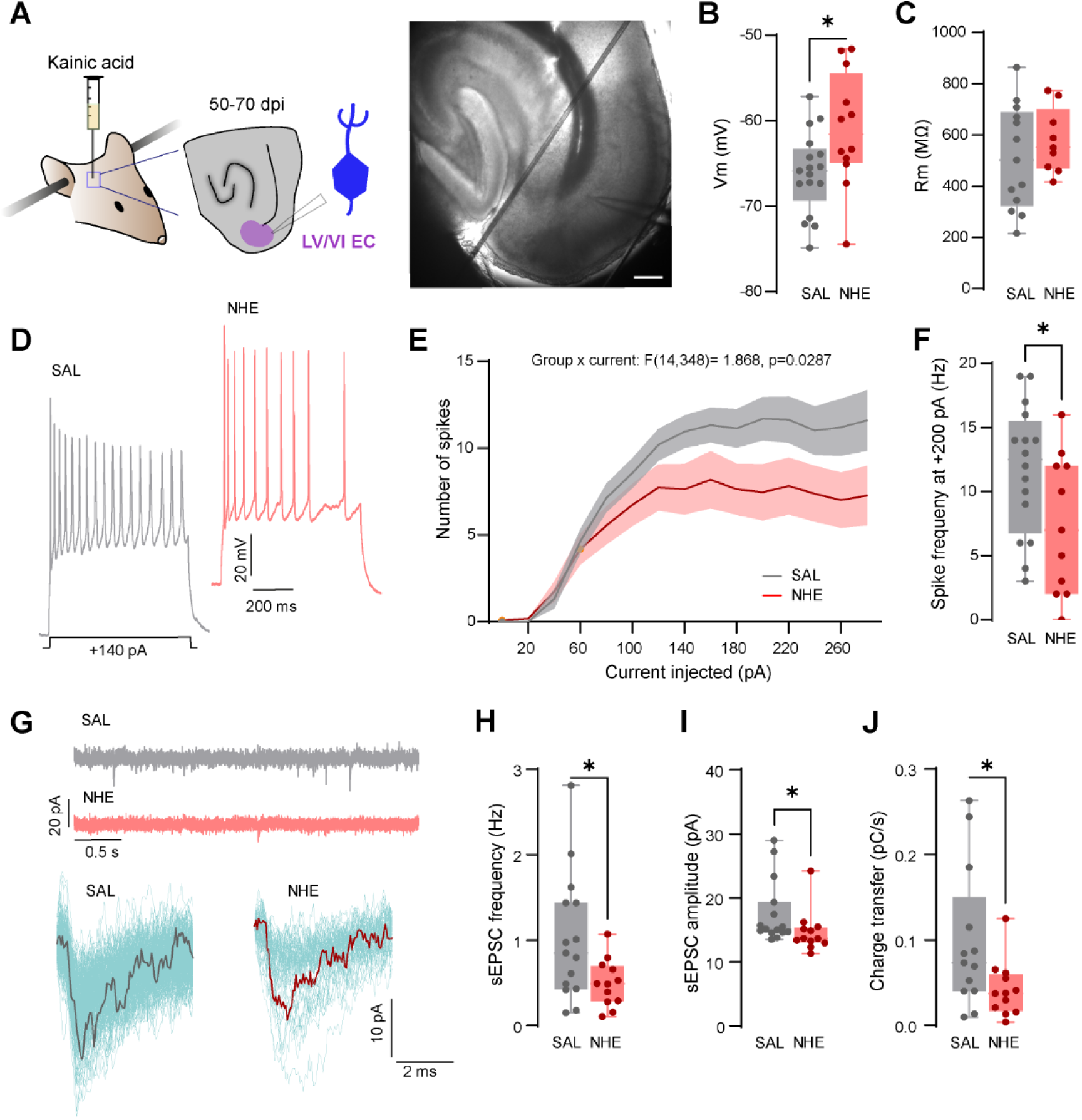
Reduced evoked firing and excitatory synaptic drive in deep-layer entorhinal neurons. (A) Left, experimental design for whole-cell patch-clamp recordings of putative pyramidal neurons in layers V-VI of the entorhinal cortex (EC) in acute sagittal slices from SAL and NHE mice at 50-70 days post-injection (dpi). Right, transmitted-light image of a slice with the recording pipette. (B, C) Resting membrane potential (Vm; B) and input resistance (Rm; C) of deep-layer EC neurons. (D) Representative voltage responses to a depolarizing current step (+140 pA). (E) Input-output curves showing the number of action potentials evoked by increasing current injections (group × current interaction: F(14,348) = 1.868, p = 0.0287). (F) Action potential firing frequency at +200 pA. (G) Top, representative recordings of spontaneous excitatory postsynaptic currents (sEPSCs) held at −70 mV. Bottom, superimposed individual events (cyan) and averaged waveform (dark line) from a representative neuron of each group. (H-J) sEPSC frequency (H), amplitude (I) and charge transfer (J). SAL n = 15 cells from 3 mice, NHE n = 12 cells from 4 mice. Unpaired t-test or Mann-Whitney test for B, C, F and H-J; two-way repeated-measures ANOVA or mixed-effects model for E. Data are shown as box-and-whisker plots (median, interquartile range and minimum-maximum) with individual cells overlaid, and as mean ± SEM (shaded) for the input-output curves. *p < 0.05. Scale bar: 100 µm

Analysis of passive membrane properties revealed a significantly more depolarized resting membrane potential in NHE neurons, whereas input resistance remained comparable between groups (Fig. 4B-C). Despite this membrane depolarization, NHE neurons exhibited reduced action potential firing in response to depolarizing current injections, with a lower firing response particularly evident at higher current intensities (Fig. 4D-E). Consistent with this observation, the number of action potentials elicited by a +200 pA current injection was significantly reduced in NHE neurons (Fig. 4F).

To further characterize the functional alterations observed in deep-layer EC neurons, we next examined their spontaneous excitatory synaptic activity. NHE neurons exhibited a significant reduction in spontaneous excitatory postsynaptic current (sEPSC) frequency, accompanied by decreased sEPSC amplitude and charge transfer (Fig. 4H-J).

Together, these findings reveal persistent alterations in both intrinsic firing responses and excitatory synaptic activity in deep-layer EC neurons following NHE. Notably, these single-cell alterations coexist with a reduced proportion of active electrodes in EC, despite the absence of significant differences in overall spontaneous firing and bursting frequency at the network level.

### Region-specific changes in the network response to a pharmacological challenge

To assess whether the altered network activity was accompanied by changes in how the circuit responds to a challenge, we applied the kainate receptor agonist ATPA (10 µM), which preferentially activates GluK1-containing receptors, and compared burst frequency in each region before and during ATPA application in SAL and NHE slices (Fig. 5A-C).

**Figure 5.**
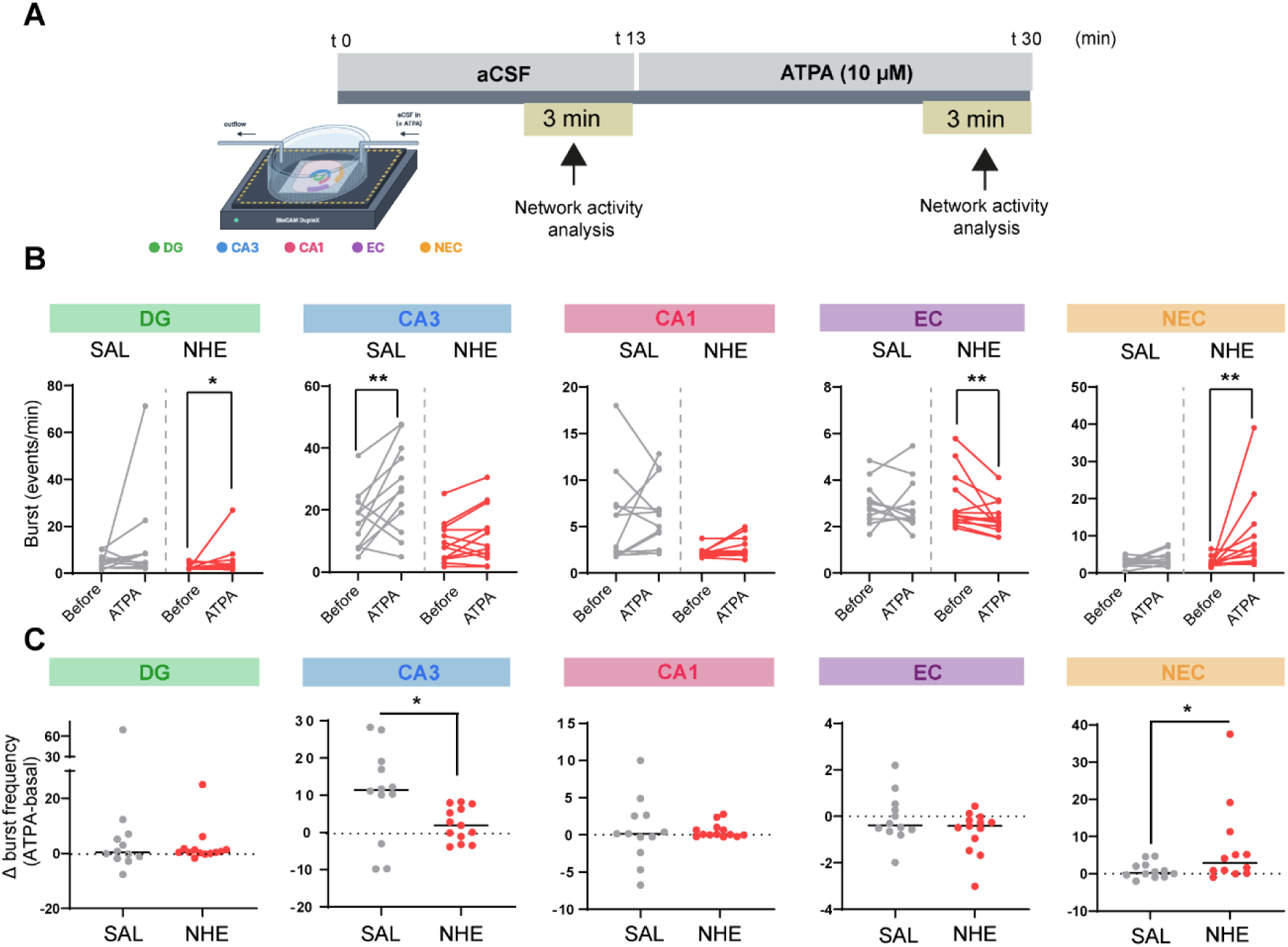
Region-specific changes in the response to a pharmacological challenge. (A) (A) Experimental protocol. Slices on HD-MEAs were perfused with aCSF (0-13 min) and then with the GluK1 kainate receptor agonist ATPA (10 µM; 13-30 min). Network activity was analyzed in the last 3 min of each condition. (B) Burst frequency before and after ATPA (10 µM) in DG, CA3, CA1, EC and NEC slices from SAL and NHE mice. (C) Change in burst frequency (ATPA − baseline) per slice. For DG, CA3, CA1 and EC: SAL n = 12 slices from 3 mice, NHE n = 13 slices from 3 mice; for NEC: SAL n = 12 slices from 3 mice, NHE n = 12 slices from 3 mice. Wilcoxon matched-pairs signed-rank test for CA3, EC and NHE in NEC, and paired t-test for SAL in NEC (B). Unpaired t-test for CA3 and Mann-Whitney test for NEC (C). Each dot represents one slice. Lines connect paired measurements from the same slice (B), and horizontal lines indicate the median (C). *p < 0.05, **p < 0.01.

ATPA modified burst frequency in a region- and group-dependent manner. In SAL slices, ATPA increased bursting in CA3 but not in the other regions. In NHE slices, ATPA increased burst frequency in DG and NEC, and decreased it in EC, whereas CA3 and CA1 were unaffected (Fig. 5B).

To compare responses between groups, we quantified the change in burst frequency induced by ATPA (ATPA-baseline) in each slice (Fig. 5C). The change did not differ between SAL and NHE in DG, CA1 or EC. In contrast, the response was significantly attenuated in CA3 and significantly enhanced in NEC in NHE compared with SAL slices. The decrease in bursting observed in EC of NHE slices at the within-group level, however, did not translate into a difference between groups.

Together, these results indicate that NHE alters the network response to ATPA in only two regions, and in opposite directions: attenuated in CA3 and enhanced in NEC.

## Discussion

Here we show that a single injection of low-concentration kainic acid, which induces NHE without seizures, leaves a persistent and heterogeneous footprint in the hippocampal-cortical circuit two months later. Ex vivo, the network was hypoactive: firing and bursting were reduced in the DG, CA3 and CA1, the proportion of active electrodes was lower in all recorded regions, and the directional coupling from CA1 to the EC was weakened. At the cellular level, however, there was no sign of a general loss of excitability. CA1 pyramidal neurons showed higher input resistance and increased evoked firing, CA3 pyramidal neurons fired more at higher current intensities, CA3 interneurons were hyperexcitable, and deep EC neurons received fewer spontaneous excitatory inputs. This study shows that cellular and network remodeling are dissociated, as, in contrast to the common assumption, hyperexcitable neurons do not necessarily translate into a hyperactive network.

The cellular changes were not uniform across cell types or regions, which is itself informative of regional selectivity. CA1 pyramidal neurons became more excitable, with higher input resistance and a reduced hyperpolarization-activated inward current ^38^, consistent with a lasting increase in intrinsic excitability after the insult, as described after other forms of injury ^8,9,11,12^. In CA3, pyramidal neurons required more current to fire but fired more at higher intensities, while interneurons fired more from intermediate current intensities onward. This enhanced firing was accompanied by altered voltage-dependent currents in response to hyperpolarization, including a reduced hyperpolarization-activated inward current, a conductance that shapes how CA3 interneurons integrate their inputs ^39^. Whether this change contributes to their hyperexcitability remains to be tested. In contrast, CA1 interneurons showed a marked reduction in repetitive firing together with a shorter latency to the first spike, a pattern that could reflect an early loss of firing capacity, such as depolarization block, although our recordings do not resolve it. Alterations in interneuron firing are well described in epilepsy and in Alzheimer’s disease models ^40–45^, and changes in the delay to the first spike point to altered subthreshold potassium conductances ^46,47^.Deep EC neurons were more depolarized yet fired less, a combination that is also compatible with reduced excitability from a depolarized state ^22,23^. Excitability measured in single neurons thus does not predict how a region behaves within the circuit, in line with work showing that neurons with altered synaptic properties can lack somatic hyperexcitability ^15^ and that opposite regional effects can coexist after a kainate insult ^13^.

Several, non-exclusive mechanisms could explain how hyperexcitable neurons coexist with a hypoactive network. First, in CA3, the increased firing of interneurons may enhance feedforward and feedback inhibition onto principal cells and reduce network output even if pyramidal neurons are intrinsically more excitable ^25,43^. Second, the reduced activity of the DG and CA3 may decrease the drive onto CA1, whose pyramidal neurons are more excitable but depend on that input to fire spontaneously in the slice. Third, homeostatic processes that operate at the circuit level may have counterbalanced the initial hyperexcitability and, over time, overshot toward lower activity ^16–19^. A further possibility relates to what the HD-MEA measures. Extracellular spikes were not assigned to cell types ^48^, so the reduced firing in each region may be driven unevenly by principal neurons or interneurons. In CA1, where interneurons fired less in response to current injection while pyramidal neurons fired more, the net reduction in spontaneous activity could partly reflect the contribution of interneuron spikes, which are often fired at high rates ^49^. This would not apply to CA3, where interneurons fired more, and it would be expected to disinhibit pyramidal neurons rather than silence them, so it cannot by itself account for the network hypoactivity. Finally, the reduced excitatory synaptic drive onto deep EC neurons, together with the weakened CA1-to-EC coupling, suggests that the output of the hippocampal circuit is selectively impaired. We did not test these possibilities directly, and they should be taken as working hypotheses. That the direction of activity changes after an insult can depend on the time elapsed, with an early hypoactivity that later turns into hyperactivity, has been reported in cortex after trauma ^14^.

The EC illustrates how network and cellular readouts can differ. Its spontaneous firing and bursting did not change, yet the proportion of active electrodes was lower, and single neurons were less excitable and received less excitatory input. Because firing rate was computed on active electrodes, these observations are compatible with a fraction of units that became silent while the remaining ones kept their rates. The same reasoning applies to the other regions, where the lower proportion of active electrodes may reflect silenced neurons and, potentially, neuronal loss, which we did not quantify.

The response to ATPA indicates that the network is not simply silenced but reorganized. The effect of ATPA differed between saline and NHE groups only in CA3, where it was attenuated, and in the non-entorhinal cortex, where it was enhanced, with a difference in firing only in the latter. The other regions responded similarly in both groups. This regional pattern argues against a global suppression and suggests that the capacity of the circuit to respond to a kainate-receptor agonist is selectively redistributed. We cannot attribute the ATPA effect to a specific receptor subunit, although it preferentially activates GluK1-containing receptors, which regulate inhibitory transmission in the hippocampus ^31^ and are involved in kainate-driven network activity ^50^, and our design was paired and lacked a vehicle control, so these results describe a differential response rather than a defined mechanism.

An important point is the apparent discrepancy between our ex vivo hypoactivity and the hyperexcitability observed in vivo. In this model, in vivo recordings show epileptiform spikes up to 50 days after the injection ^26^. These observations are not necessarily contradictory. Acute slices are deafferented and lack the extrinsic and neuromodulatory inputs that sustain activity in vivo; our measures capture mean firing rate and bursting in spontaneous activity, whereas in vivo epileptiform events reflect synchronous population discharges that may depend on connectivity that is cut in the slice; and the vulnerability of different cell populations to the slicing procedure may differ between groups. We therefore interpret our results as a description of the circuit’s baseline state in the absence of external drive, not as evidence that the network is hypoactive in the living animal.

Our study has limitations. Cell loss was not quantified, so we cannot rule out that part of the reduced activity reflects neuronal loss, although tissue markers did not converge with those of epilepsy ^27,28^; sections are available for this analysis. Patch-clamp and HD-MEA recordings were made in different slice preparations (sagittal and horizontal, respectively), so cellular and network measures are not from the same tissue. The HD-MEA does not identify the cell type of the recorded spikes, so we cannot determine whether the changes in network activity arise from principal neurons, interneurons, or both. We examined a single time point and an ex vivo preparation, and we did not address the temporal evolution of these changes.

In summary, a hyperexcitability insult without seizures can leave a lasting, region-specific remodeling in which cellular and network properties no longer move together. Considering only the cell or only the network would have given an incomplete, and in part opposite, picture.

Because hyperexcitability without seizures is also found in neurodegenerative conditions such as Alzheimer’s disease ^1–4^, a dissociation between cellular and network properties like the one described here could be relevant to understand circuit dysfunction in those contexts ^41,44,45^, a possibility that will require testing in appropriate models. Future work should assign HD-MEA spikes to cell types, for example by waveform features ^48,51^, resolve the time course of this dissociation, define the contribution of interneuron subtypes, and combine in vivo and ex vivo recordings to relate the baseline state of the circuit to its activity in the living animal.

## Author contributions

**D.V.H.:** Investigation, Methodology, Formal analysis, Visualization, Writing - original draft**; J.D.O.O.:** Investigation, Methodology, Formal analysis, Visualization, Writing - original draft**; L.B.A.:**Investigation, Formal analysis; **M.A.:** Methodology, Formal analysis, Visualization, Writing - original draft; **J.M.E.-P.:** Supervision, Funding acquisition, Visualization, Writing - original draft**; N.L.:** Conceptualization, Methodology, Investigation, Formal analysis, Supervision, Funding acquisition, Writing - review & editing..

All authors reviewed and approved the final version of the manuscript.

## Supplementary figures

**Supplementary Figure 1.**
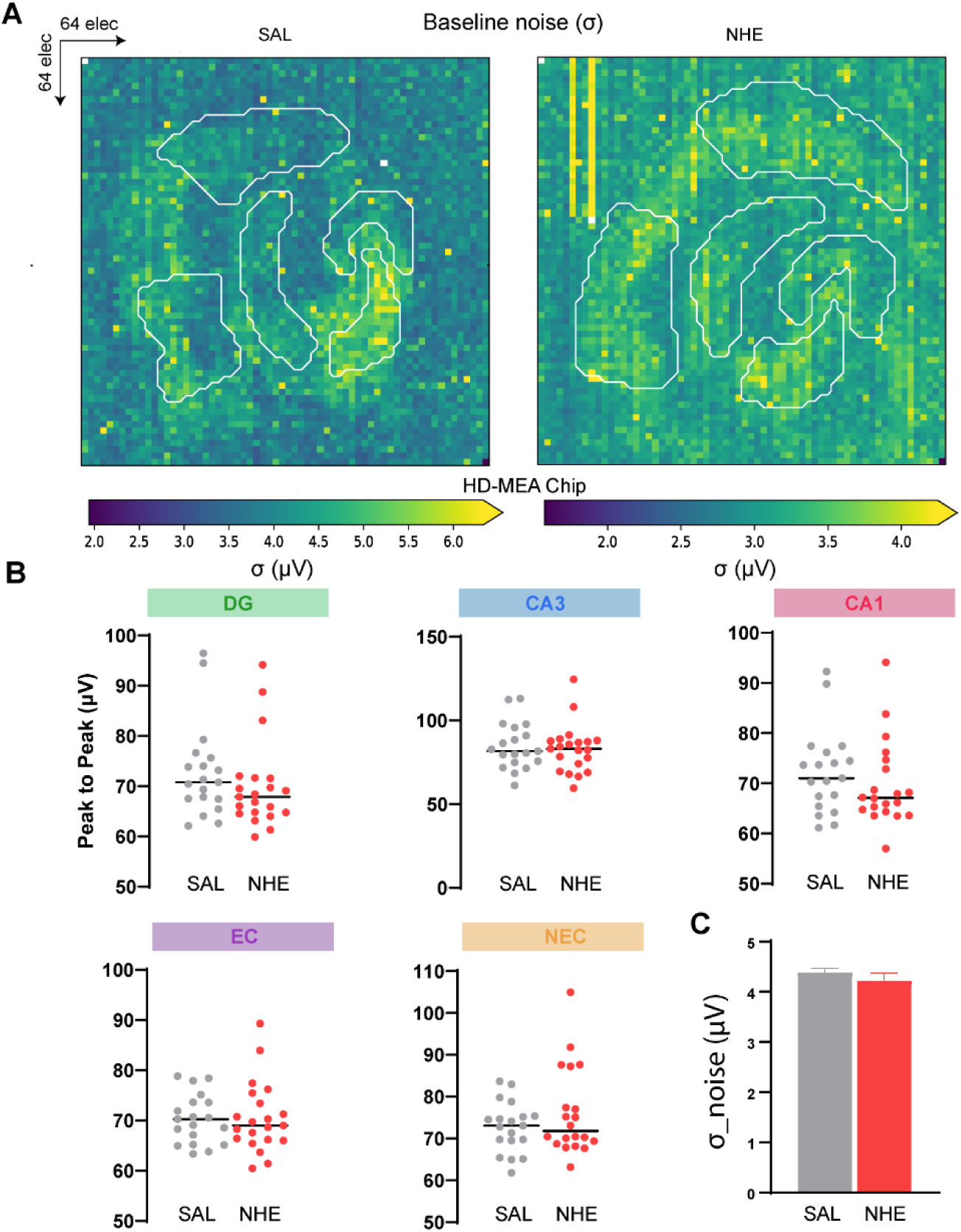
Recording quality is comparable between SAL and NHE slices. Representative maps of baseline noise (σ) across the 64 × 64 electrode HD-MEA chip for SAL and NHE slices. White outlines delineate the regions of interest. Note the different color scales. (B) Peak-to-peak amplitude of detected spikes in each region (DG, CA3, CA1, EC and NEC). (C) Mean baseline noise (σ) per chip. SAL n = 19 slices from 5 mice, NHE n = 21 slices from 5 mice. Each dot represents one slice, and horizontal lines indicate the median (B). Bars represent the mean ± SEM (C).

**Supplementary Figure 2.**
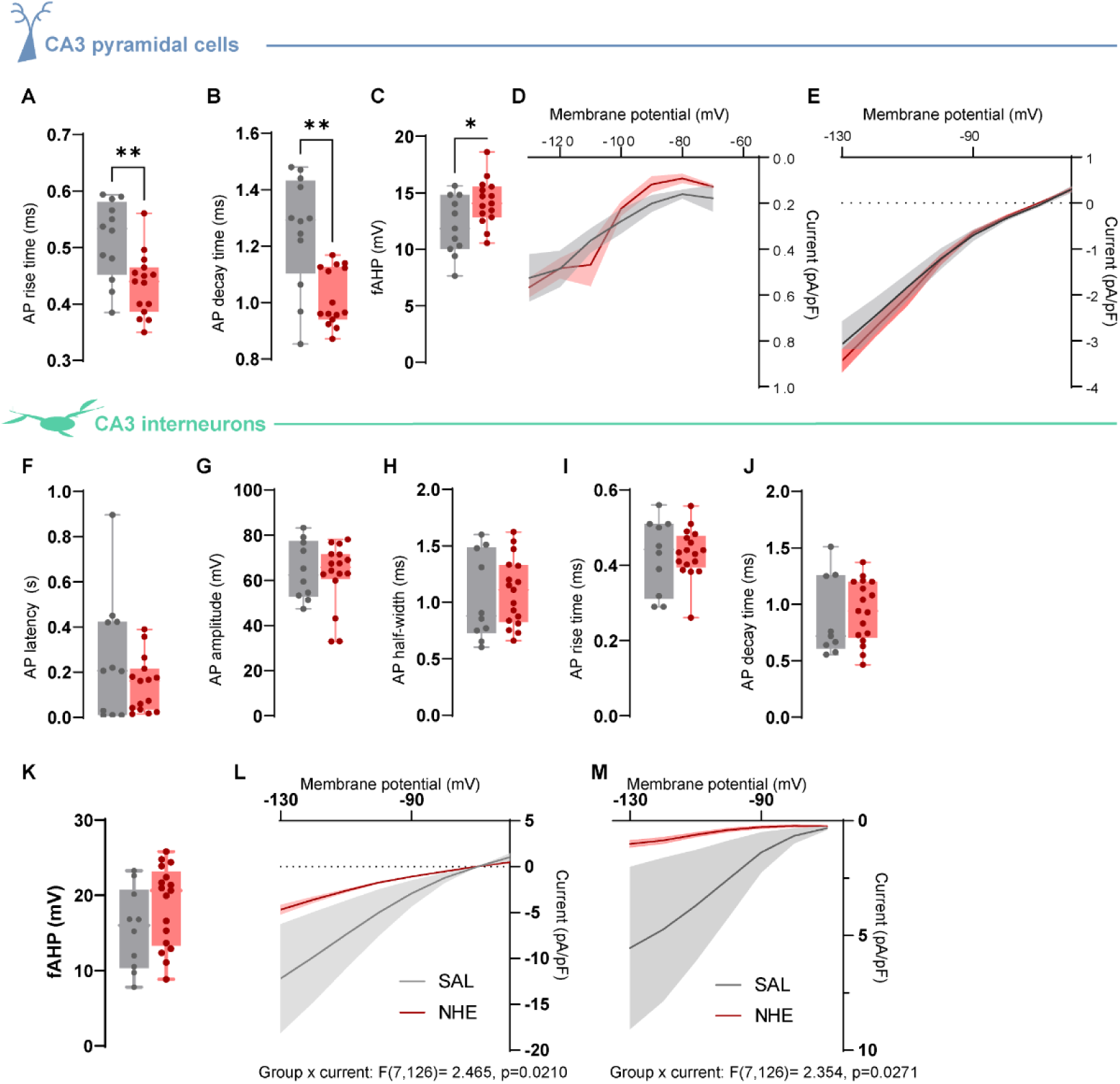
Faster action potential kinetics in CA3 pyramidal cells and a reduced hyperpolarization-activated current in CA3 interneurons at 50-70 dpi. (A-E) CA3 pyramidal neurons: action potential (AP) rise time (A), AP decay time (B), fAHP amplitude (C), hyperpolarization-activated inward current density (D) and steady-state current-voltage relationship (E). (F-M) CA3 interneurons (Dlx+): AP latency (F), AP amplitude (G), AP half-width (H), AP rise time (I), AP decay time (J), fAHP amplitude (K), steady-state current-voltage relationship (L) and hyperpolarization-activated inward current density (M). Pyramidal neurons: SAL n = 11 cells from 5 mice, NHE n = 15 cells from 5 mice; interneurons: SAL n = 12 cells from 6 mice, NHE n = 18 cells from 9 mice. Unpaired t-test for A-C. Two-way repeated-measures ANOVA, significant group × voltage interaction (L, M). Data are shown as box-and-whisker plots (median, interquartile range and minimum-maximum), with individual cells overlaid, and as mean ± SEM (shaded) for current-voltage curves. *p < 0.05, **p < 0.01.

